# Detection of pathogenic bacteria in ticks from Isiolo and Kwale counties of Kenya using metagenomics

**DOI:** 10.1101/2023.12.21.572930

**Authors:** Bryson Brian Kimemia, Lillian Musila, Solomon Lang’at, Erick Odoyo, Stephanie Cinkovich, David Abuom, Santos Yalwala, Samoel Khamadi, Jaree Johnson, Eric Garges, Elly Ojwang, Fredrick Eyase

**Affiliations:** Department of Emerging Infectious Diseases, United States Army Medical Research Directorate-Africa (USAMRD-A); Centre for Virus Research, Kenya Medical Research Institute (KEMRI), Nairobi, Kenya; Centre for Microbiology Research, Kenya Medical Research Institute (KEMRI), Nairobi, Kenya; Jomo Kenyatta University of Agriculture and Technology (JKUAT), Nairobi, Kenya; United States Armed Forces Health Surveillance Division, Global Emerging Infections Surveillance Branch, Silver Spring, Maryland, United States; United States Armed Forces Pest Management Board, United States, Silver Spring, Maryland, United States

## Abstract

Ticks are arachnid ectoparasites which rank second only to mosquitoes in the transmission of human diseases including bacteria responsible for anaplasmosis, ehrlichiosis, spotted fevers, and Lyme disease among other febrile illnesses. Due to paucity of data on bacteria transmitted by ticks in Kenya, this study undertook a bacterial metagenomic-based characterization of ticks collected from Isiolo, a semi-arid pastoralist County in Eastern Kenya, and Kwale, a coastal County with monsoon climate on the southern Kenyan border with Tanzania. A total of 2,918 ticks belonging to 3 genera and 10 species were pooled and screened in this study. Tick identification was confirmed through the sequencing of Cytochrome C Oxidase Subunit 1 (COI) gene. Bacterial 16S rRNA gene PCR amplicons obtained from the above samples were sequenced using the MinION (Oxford Nanopore Technologies) platform. The resulting reads were demultiplexed in Porechop, followed by trimming and filtering in Trimmomatic before clustering using Qiime2-VSearch. A SILVA database pretrained naïve Bayes classifier was used to taxonomically classify the Operational Taxonomic Units (OTUs). The bacteria of clinical interest detected in pooled tick assays were as follows: *Rickettsia spp.* 59.43% of pools, *Coxiella burnetii* 37.88%, *Proteus mirabilis* 5.08%, *Cutibacterium acnes* 6.08% and *Corynebacterium ulcerans* 2.43%. These bacteria are responsible for spotted fevers, query fever (Q-fever), urinary tract infections, skin and soft tissue infections, eye infections and diphtheria-like infections in humans, respectively. *P. mirabilis, C. acnes* and *C. ulcerans* were detected only in Isiolo. Additionally, COI sequences allowed for the identification of *Rickettsia* and *Coxiella* species to strain level in some of the pools. Diversity analysis revealed that the tick genera had high levels of Alpha diversity but the differences between the microbiomes of the three tick genera studied were not significant. The detection of *Cutibacterium acnes*, commonly associated with human skin flora suggests that the ticks may have contact with humans potentially exposing them to bacterial infections. The findings in this study highlight the need for further investigation into the viability of these bacteria and the competency of ticks to transmit them. Clinicians in these high-risk areas also need to be appraised for them to include Rickettsial diseases and Q-fever as part of their differential diagnosis.

## Introduction

Interaction between humans and domestic animals creates a pathway for ectoparasites, such as ticks, leading to the emergence of zoonotic diseases—an issue of increasing concern according to the World Health Organization’s One Health concept (1–4). Among these diseases, tick-borne bacterial infections like Lyme disease, anaplasmosis, ehrlichiosis, and rickettsiosis pose significant global health threats (5). These diseases not only cause debilitating symptoms but, if left untreated, can result in chronic health issues and fatalities, imposing a substantial burden on healthcare systems and compromising a country’s economy (5,6).

Despite the global prevalence of tick-borne pathogens, accurate diagnostics are limited, potentially under-reporting their actual occurrence (7,8). In regions like Kenya, with diverse tick fauna and abundant livestock and wildlife populations, Lyme disease, anaplasmosis, and rickettsial diseases have been reported with varying prevalence (9,10). Spotted Fever Group *Rickettsias* such as Rocky Mountain Spotted Fever, Rickettsial-pox, and Boutonneuse fever have been reported in North and South America, Europe, Africa, and Asia. In fact, the most reported spotted fever reported in the United States is African Tick Bite Fever, caused by *Rickettsia africae* infection after international travel from Africa to the United States(11) Continuous surveillance and research are vital to understanding the distribution and impact of these diseases, crucial for developing effective prevention and control strategies (10).

This study addresses this gap by focusing on two specific regions in Kenya: Isiolo and Kwale counties. Isiolo is an arid/semi-arid county where livestock keeping constitutes the primary income source for approximately 80% of its population (12). Hosting a large livestock market that attracts sellers from surrounding counties (13), it represents a critical hotspot for understanding zoonotic disease dynamics. Kwale, primarily an agricultural county, is home to a significant livestock-keeping community, especially in the Nyika plateau region (14). Kwale county also borders Tanzania, this provides a cross-border importation route for vectors and by extension pathogens from the southern border. These unique economic activities and environmental characteristics make both Isiolo and Kwale ideal study areas, offering valuable insights into the prevalence and diversity of tick-borne bacterial diseases in distinct contexts.

By utilizing 16S rRNA gene sequencing, this research aims to comprehensively analyse the microbiomes of ticks collected from these regions. The study’s objectives include assessing bacterial diversity through metagenomic analysis, the identification of potential and novel pathogenic bacteria, and confirming the taxonomic identity of tick species carrying tick-borne bacteria through metabarcoding. This focused approach ensures a nuanced understanding of the prevalence and nature of tick-borne diseases, essential for tailoring targeted public health management and prevention strategies in these areas.

## Materials and Methods

### Ethical Approval

The study was approved by the Kenya Medical Research Institute (KEMRI) Scientific and Ethics Review Unit (SERU) under protocol number KEMRI/SERU/CCR/4431. The study was carried out under National Commission for Science, Technology & Innovation (NACOSTI) license 421618. The study was submitted for approval to the Walter Reed Army Institute of Research (WRAIR) Human Subjects Protection Branch (HSPB) and Institutional Review Board (IRB) under package WRAIR #3000. The study was exempted from requiring any further approval as per WRAIR policy #25 because no samples, data or information was being collected from human subjects. Consent was sought from farmers and pastoralists using an informed consent form.

### Sampling and Pooling

Ticks were collected from cattle, sheep, goats, and camels from Isiolo and Kwale counties. Sites from Isiolo County were Merti, Shambole, Isiolo market and Isiolo slaughterhouse (Fig 1a). The ticks were collected by veterinarian through grooming. Two sites were sampled in Kwale County, Kisima and Mlalani (Fig 1b). Sampled ticks were placed in 15ml centrifuge tubes (Corning Inc, Corning, NY, USA) and transported under dry ice to the KEMRI/ United States Army Medical Research Directorate – Africa (USAMRD-A) laboratories. After reception, the ticks were identified morphologically under a microscope using taxonomic keys (15–19) and pooled in groups of 1 to 8 individuals based on species, size, and site of collection before transferring to cryovials for storage at -81°C.

**Fig 1:**
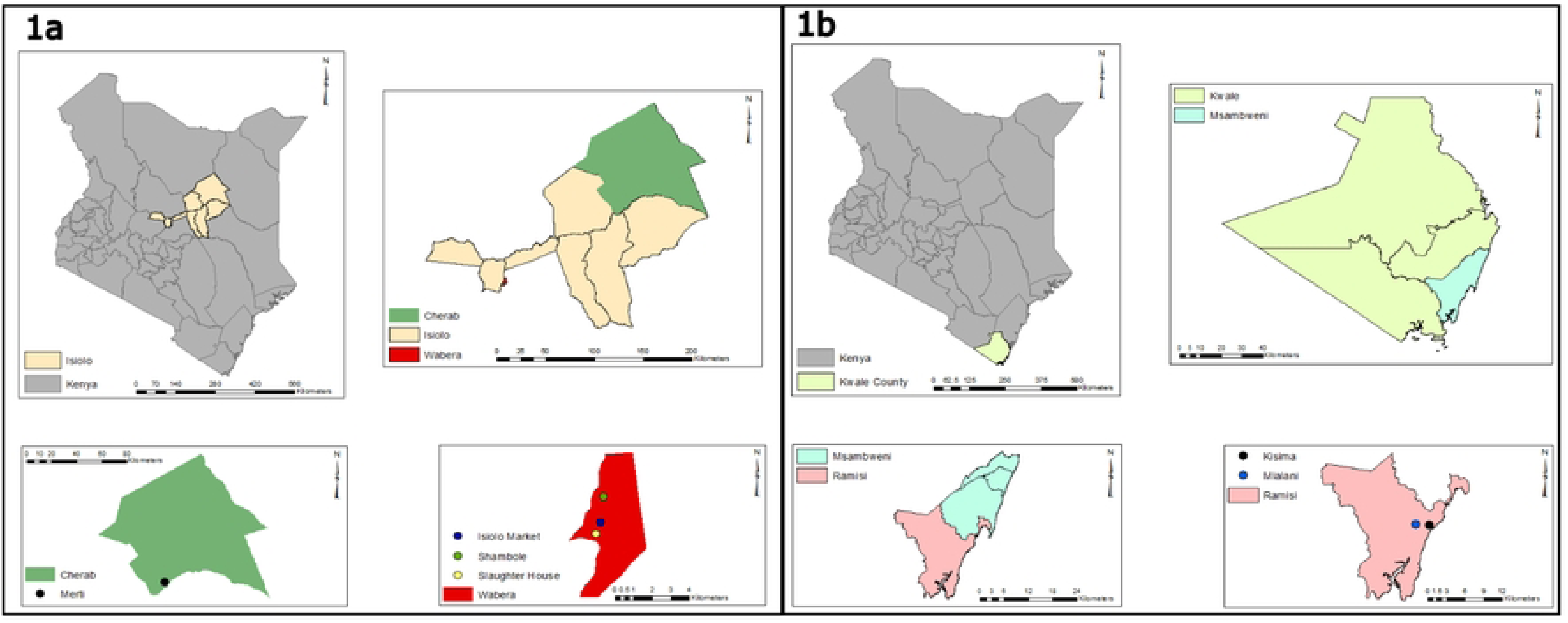
Map of Sampling Sites, Isiolo County (a) Kwale County (b).

### Surface Sterilization and Homogenization

To ensure removal of external bacterial contaminants, surface sterilization was performed on the ticks. Briefly, cryovials were placed on ice in a biosafety cabinet and 1ml of 5% Sodium Hypochlorite solution was dispensed into the open cryovials and allowed a contact time of 5 mins. The ticks were then rinsed 3 times with distilled water.

Following surface sterilization, homogenization was carried out as follows: the ticks, were frozen at -81°C for a minimum of 30 minutes or exposed to liquid nitrogen in tubes with bashing beads to render them brittle. The ticks were then homogenized using an Omni bead Ruptor 24 (Omni International, Inc, Kennesaw, GA, USA) for two cycles, running at 4m/s for 60 seconds with a 10-second pause between cycles. 1000µl of homogenization media (40% glycerol, without antibiotics/antimycotic) was added followed by pulse vortexing to mix. The homogenate was then centrifuged at 10,000 rpm for 10 minutes. Subsequently, 200µl of the supernatant was withdrawn for DNA extraction, while the remainder of the homogenate was stored at -81°C for future use.

### DNA Extraction and PCR Amplification

DNA extraction was performed using the Quick-DNA™ Fungal/Bacterial Miniprep Kit (Zymo Research, Orange, CA) per the manufacturer’s instructions. Two separate PCR amplifications were performed to amplify the 16s rRNA and COI genes. The first was for bacterial metagenomics while the second was to confirm the morphological tick species identification. For 16S rRNA PCR amplification, 10 µM of 27F (5′-AGAGTTTGATCCTGGCTCAG-3′) forward primer and 10 µM of 1492R (5′-TACGGYTACCTTGTTACGACTT-3′) (20) reverse primer pair used whereas for the Cytochrome Oxidase Subunit 1 (COI), 10µM of LCO1490 (5’-GGTCAACAAATCATAAAGATATTGG-3’) forward primer and 10 µM of HC02198 (5’-TAAACTTCAGGGTGACCAAAAAATCA-3’) (21) reverse primer pairs were used. The NEBNext High Fidelity 2x PCR Master Mix (New England Biolabs, Ipswich, MA) was used according to the manufacturer’s instructions. The PCR cycles were as follows; For 16S rRNA, 95°C initial denaturation for 5 min, 35 cycles of 95°C denaturation for 30 sec min, 53.8°C annealing for 30 sec and 72°C extension for 1.5 min and a final extension at 72°C for 7 min. The COI PCR amplification cycle was as follows; 95°C initial denaturation for 5 min, 35 cycles of 95°C denaturation for 30 sec min, 49°C annealing for 30 sec and 72°C extension for 45 sec and a final extension at 72°C for 7 min.

A 0.8%w/v TBE agarose gel was used to confirm the presence of bands at approximately 1500 base pairs (bps) size for 16S rRNA gene and 710 bps for COI gene, indicating successful amplification.

### Sequencing

After DNA extraction and PCR amplification, amplicons from both 16S rRNA and COI genes were quantified to ensure that they attained the concentration (15.58ng/µl) for the Native Barcoding Kit (SQK-NBD.112.96) (Oxford Nanopore Technologies, Oxford, United Kingdom). The sequencing was performed on a MinION MK1B (Oxford Nanopore Technologies, Oxford, United Kingdom) platform using an R9.4.1 flow-cell (Oxford Nanopore Technologies, Oxford, United Kingdom). Data acquisition was done using the MikNOW Version 22.10.7 (Oxford Nanopore Technologies, Oxford, United Kingdom) software.

### Bioinformatics Analysis

The raw sequencing reads (fast5 files) were base called using Guppy Version 6.3.8 (Oxford Nanopore Technologies, Oxford, United Kingdom) into FastQ files. Quality control checks were done using PycoQC Version 2.5.2 (22). De-multiplexing, removal of barcodes and adapters was done in Porechop Version 0.2.4 (23) after which Trimmomatic Version 0.36 (24) was used for quality filtering and trimming.

16S rRNA gene amplicon reads between 1,100 and 1,700 bps and COI reads between 600 and 850bps were selected.

16S rRNA amplicon reads were imported into QIIME2(25) for de-replication, filtered for chimeras and clustered in VSearch (26) and the Operational Taxonomic Units (OTUs) classified using a naïve Bayes classifier (27) trained on 16S rRNA data from the SILVA-138-99 database. Using controls included in the sequencing run, common laboratory contaminants present were filtered out and potential pathogenic bacteria were selected.

The COI reads were then submitted to the BugSeq metagenomic platform for the identification of both Vectors and bacteria that use oxidative phosphorylation during energy production. Taxonomic ID of the tick vectors was then confirmed on the MIDORI2 server (28) (VersionGB252) using the RDP classifier option (29).

### Data Analysis

The OTU count data was first cleaned in Microsoft Excel by removing any blank spaces and confirming the alignments of columns and rows after which they were imported into RStudio (v 2023.06.1 Build 524). Alpha diversity was measured using the Shannon and Simpson indices per tick genus at the bacterial order, genus, and species level. The Shannon index measures the richness and diversity of OTUs in the genus while the Simpson index measures the Alpha diversity that provides information regarding the overall number of total individual OTUs. The indices were tested for distribution normalcy using the Shapiro-Wilk normalcy test to determine whether one way ANOVA for normally distributed data or Kruskal-Wallis non-parametric test was appropriate for statistical significance testing between the indices. Mann-Whitney U test was used to test the differences between the median number of reads for each order as a measure of abundance. This was followed by Dunn’s multiple comparison test between each of the orders.

Beta diversity between the tick genera was determined using PERMANOVA. The similarity between the number and types of OTUs between the genera was tested and a Bray-Curtis Dissimilarity matrix generated. This matrix was then tested using 999 permutations of PERMANOVA. This allowed for model testing to determine the effect of each of the OTUs on the diversity of bacteria in all the ticks sampled at an isolated (ignoring other OTUs), individual (keeping other OTUs constant) and group (accounting for all OTUs) level.

The skewed sampling of the ticks between Isiolo and Kwale prevented any inter-county comparative analysis.

## Results

### Tick Collection

A total of 2,918 ticks were collected from Isiolo [Isiolo slaughterhouse(n=1,592), Isiolo market(n=1,019), Merti(n=246) and Shambole(n=1)] and Kwale [Mlalani(n=36) and Kisima(n=24)] counties. A vast majority of the ticks (97.94%) were collected from Isiolo and the rest from (2.06%) Kwale. These ticks were then pooled based on species and site resulting into 472 pools representing approximately 8 ticks. Aliquots from original pools were further combined into 69 super-pools representing an average of 50 ticks based on site and species. The ticks investigated in this study, covered 3 genera and 10 species (Fig 2): *Ambylomma gemma* (17.4%), *Ambylomma lepidium* (20.6%), *Ambylomma variegatum* (0.2%), *Rhipicephalus appendiculatus* (1.5%), *Rhipicephalus boophilus microplus* (2.5%), *Rhipicephalus pulchellus* (6.5%), *Hyalomma dromedarii* (10.7%), *Hyalomma truncatum* (38.5%), *Hyalomma marginatum rufipes* (1.1%) and *Hyalomma albipamartum* (0.9%) collected during the month of June 2022.

**Fig 2:**
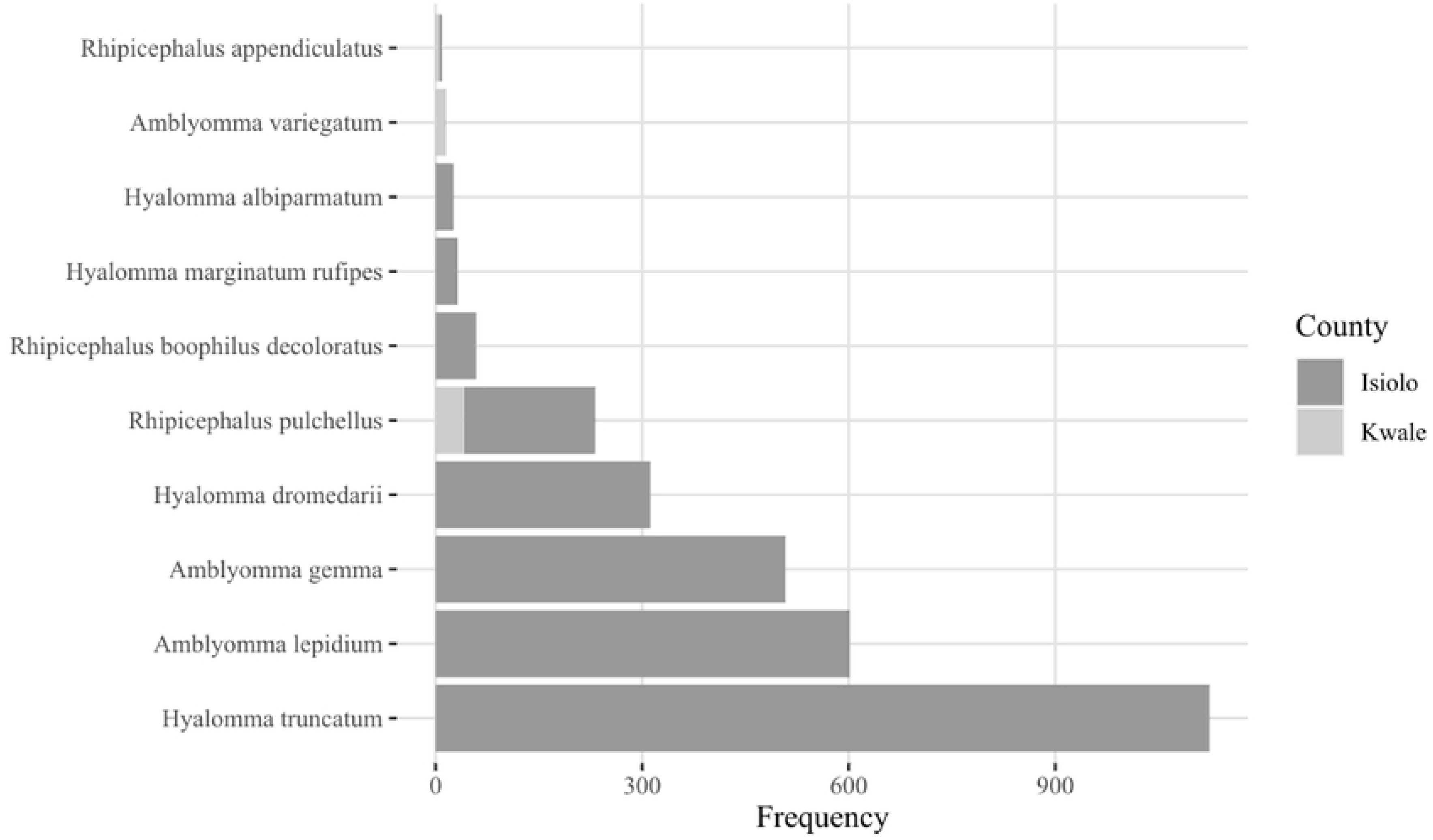
Distribution of Tick Species Collected form Isiolo and Kwale counties, individual species collected based on Cytochrome C Oxidase subunit I Identification. *Rhipicephalus boophilus microplus* was initially identified as *Rhipicephalus boophilus decoloratus* during taxonomic classification based on morphology.

### Sequencing Output

The 16S rRNA gene sequencing run generated 694.31 million bases in 547,780 reads with an estimated N50 of 1.52kb while the COI gene sequencing run generated 381.11 million bases in 430,510 reads with an estimated N50 of 845kb. This achieved the target number of reads (10,000 reads) per sample.

### Bacterial diversity and richness in different tick genera

Alpha diversity was determined using the Shannon (measure of richness) and Simpson (measure of evenness) indices (Table 1). The Shapiro-Wilkins’ test of normalcy revealed significant deviations from normal distribution of the Shannon index at the order level (p = 3.352e-08), genus level (p = 3.337e-08) and species level (p = 9.342e-05). This was also the case for the Simpson index order level (p = 2.774e-10), genus level (p = 9.923e-10) and species level (p = 1.482e-05).

**Table 1:**
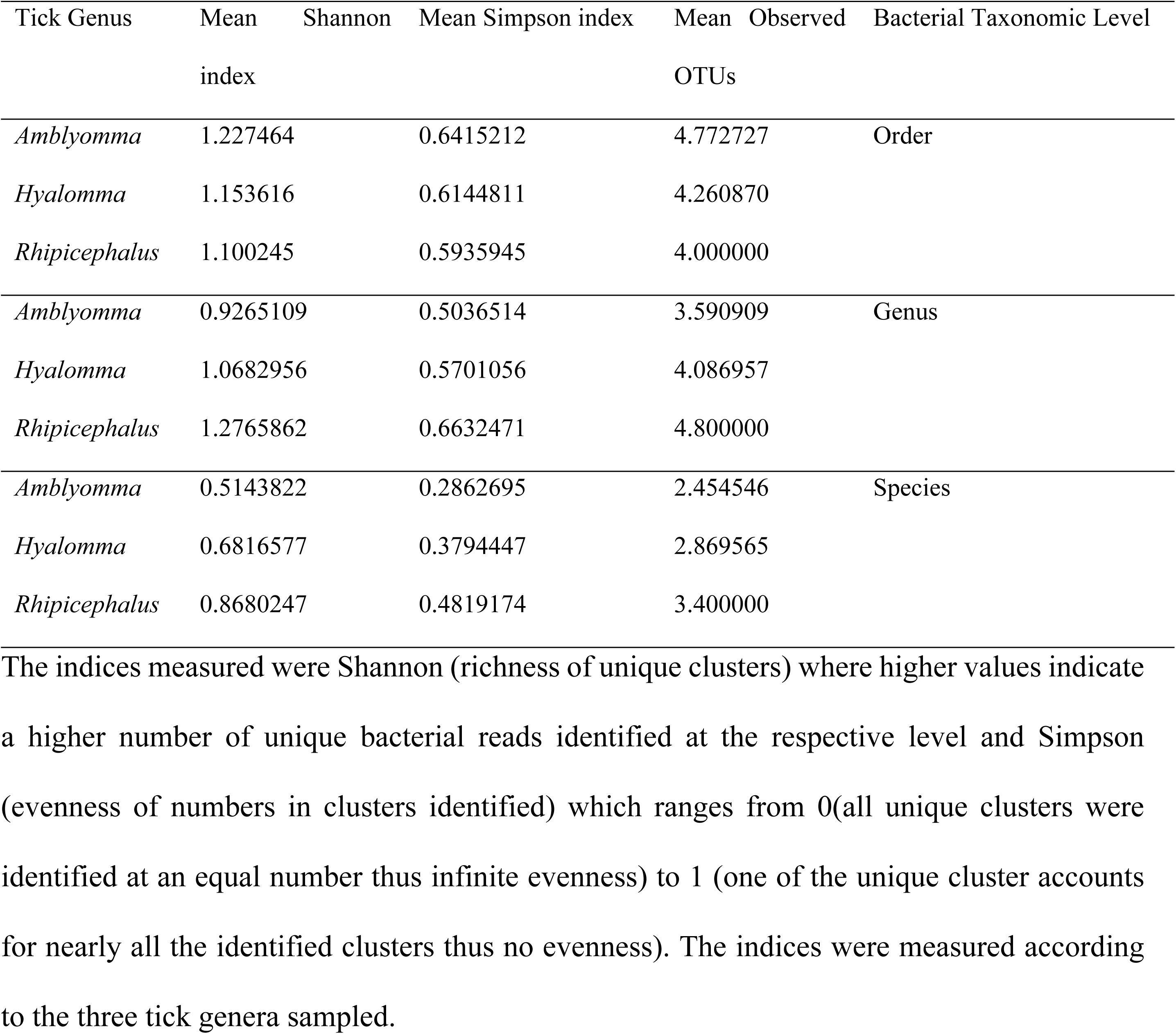
The alpha diversity of identified bacterial read clusters at the bacterial order, genus, and species level.

This informed the decision to use the Kruskal-Wallis’ test over the one-way ANOVA as the test for statistical significance between the indices at the different taxonomic levels. The results of the test showed no significant difference between the Shannon index and Simpson indices at the bacterial order, bacterial genus, and bacterial species levels (p > 0.05) when compared between the tick genera. This established that the microbiomes in the three tick genera had insignificant variations (Table 1). Comparative analysis between the two Alpha diversity indices showed that the level of richness reduces the lower the identified bacterial taxonomy level, however this drop in richness also seems to improve the evenness of the bacterial species in the compared tick genera (Fig 3). Alpha diversity analysis highlighted the big impact the *Rickettsia* genus had on the number of bacterial units identified at the bacterial genus and species level. This was because the *Rickettsia* genus could not be speciated using 16S rRNA sequences.

**Fig 3:**
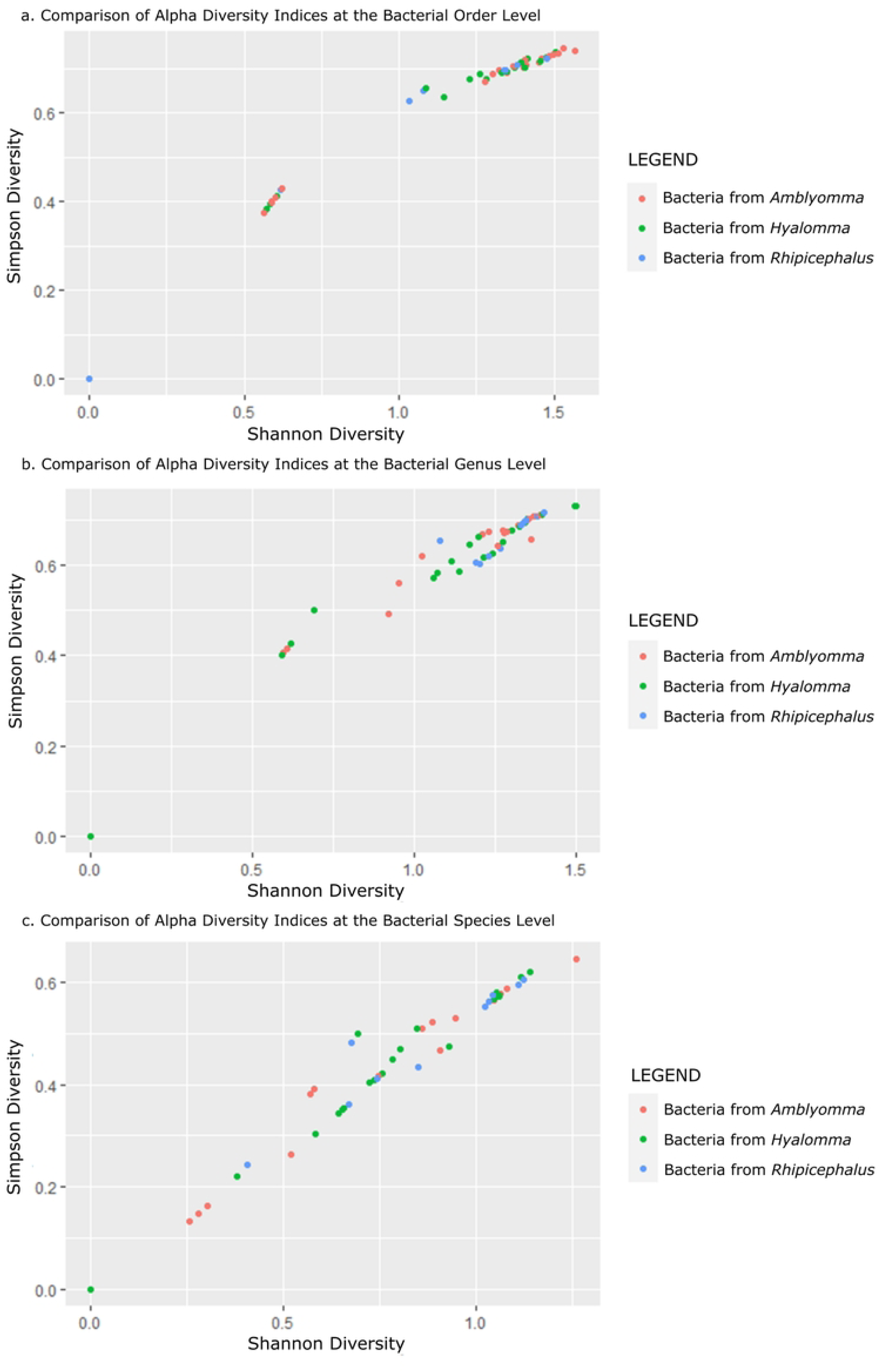
Comparative scatter plots of Alpha diversity indices of Operational Taxonomic Units at the bacterial order level (a), bacterial genus level (b) and bacterial species level (c). The three colours represent the tick genus from which they were identified from. Note the reduction in richness (Shannon diversity) and an increase in evenness (Simpson diversity) the lower the taxonomic identification level. There is marked drop in Simpson diversity between the bacterial genus and bacterial species level which coincides with the loss of the Rickettsia genus which could not be speciated.

The PERMANOVA results showed varying contribution of each of the OTUs at the isolated, individual and group level at each of the bacterial taxonomic level (Table 2). These variances were however not statistically significant (p values > 0.05).

**Table 2:**
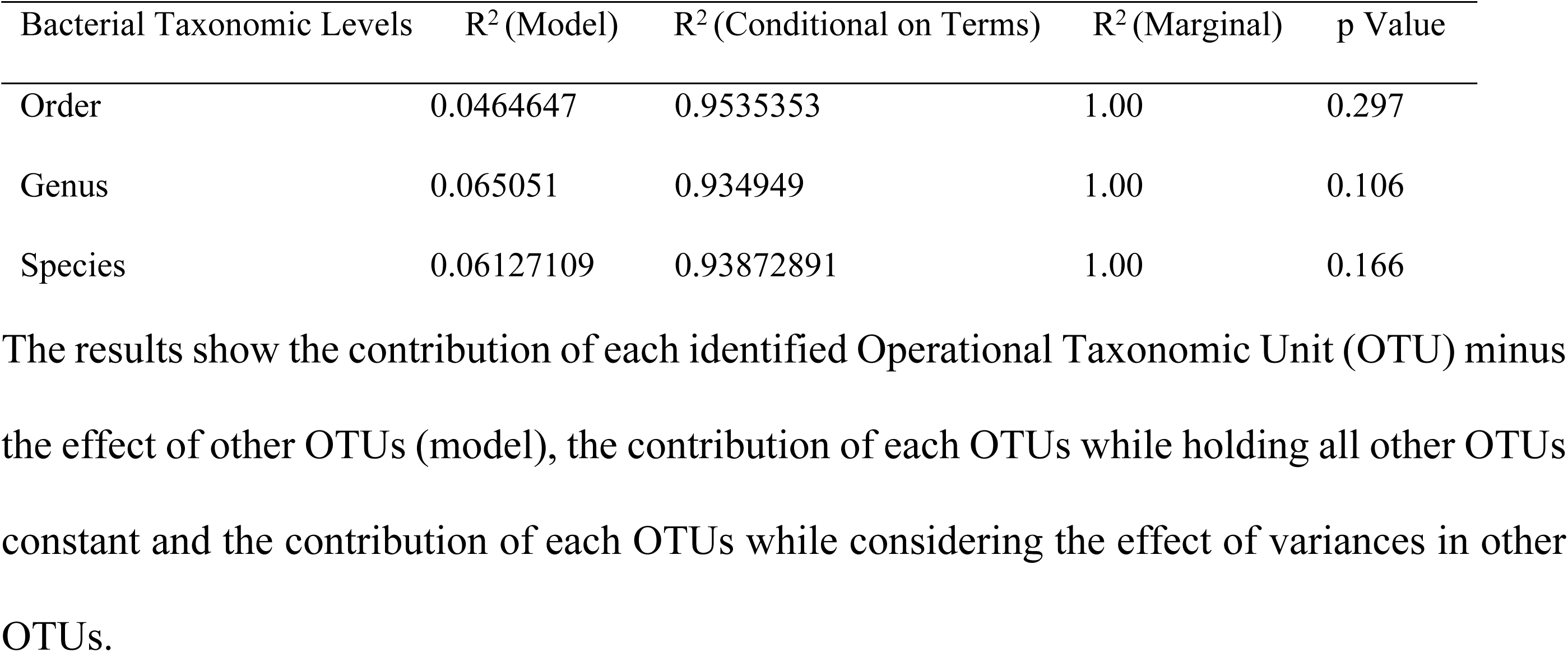
PERMANOVA Results for Beta Diversity between Tick Genera at different Bacterial Taxonomic Classification Levels.

### Taxonomic identification of bacteria and their abundances

Taxonomic read classification using QIIME2 revealed that the highest number of bacterial reads in ticks were from the order Pseudomonadales (46.52%), followed by Rickettsiales (27.53%), Coxiellales (14.57%), Staphylococcales (5.22%), Enterobacterales (4.79%), Propionibacteriales (0.54%), Entomoplasmatales (0.36%), Corynebacteriales (0.25%) and finally Micrococcales (0.23%) (Fig 4a).

**Fig 4:**
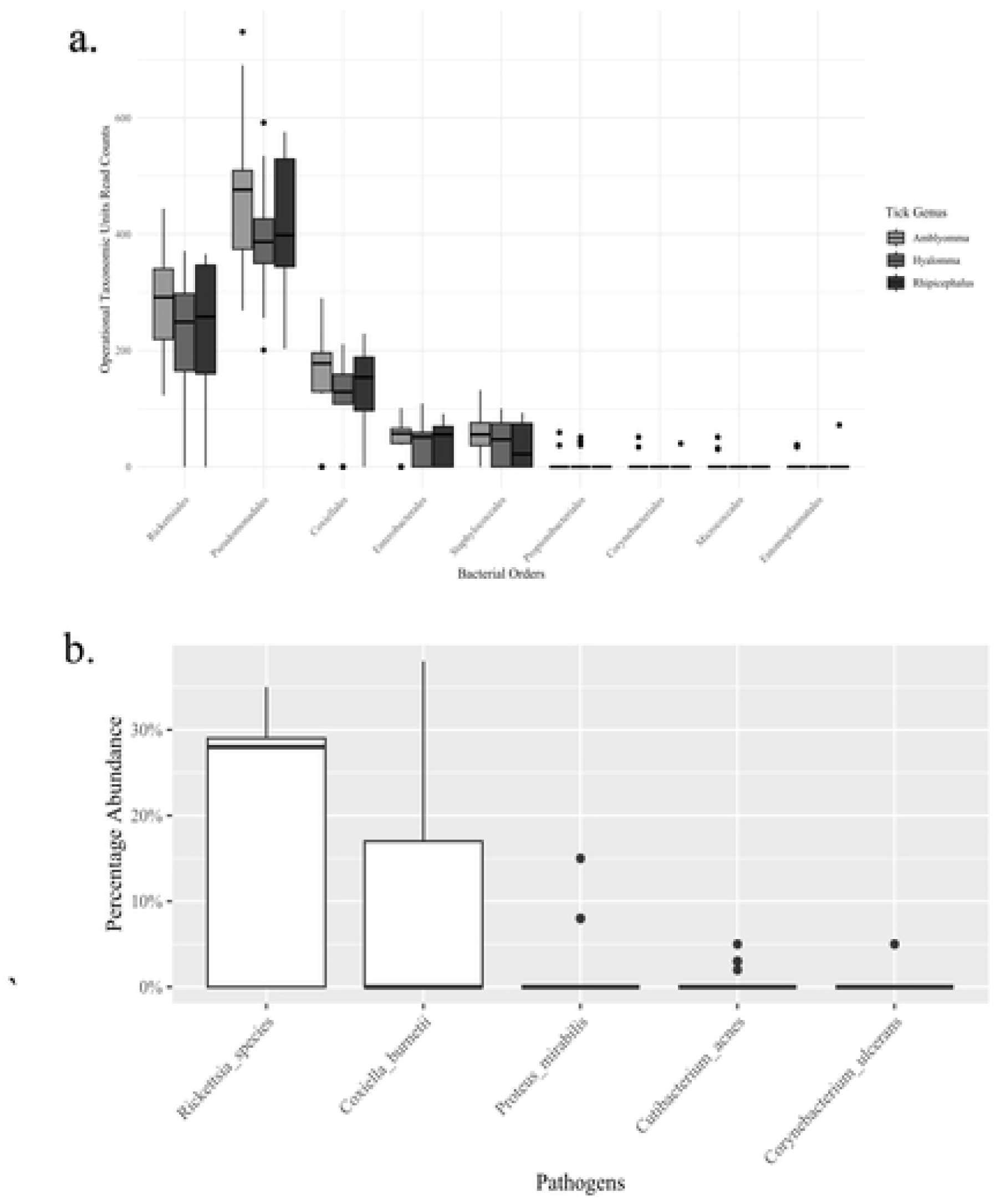
Abundances at the order level (a) and pathogenic bacteria at the genus and species levels (b). Significance between the bacterial orders was determined using Kruskal-Wallis test and yielded a p value of < 0.0001.

Kruskal-Wallis test results showed that there were significant differences (P < 0.0001) among bacterial order abundancies. However, Dunn’s multiple comparison test showed that there was no significant difference in abundancies between Rickettsiales vs coxiellales, Pseudomonadales vs Rickettsiales, Coxiellales vs Enterobacterales, Coxiellales vs Staphylococcales and Enterobacterales vs Staphylococcales. Median reads across pools for the top three orders were as follows: Pseudomonadales 410, interquartile range (IQR 353-495), Rickettsiales 252 (IQR 172-318), and Coxiellales 146 (IQR 108-187).

Comparison of the distribution of OTU reads at the order level showed that Pseudomonadales had the highest relative abundance closely followed by Rickettsiales, while the rest had a relative abundance of less than 20% (Fig 4a). For pathogenic and potentially pathogenic bacteria, the Rickettsia and Coxiella genera had the highest abundances (Fig 4b).

Six (6) bacterial OTUs were (≥60%) identified to species level, these are *Coxiella burnetii*, *Pseudomonas fluorescens*, *Proteus mirabilis*, *Staphylococcus agnetis*, *Corynebacterium ulcerans* and *Cutibacterium acnes* (Table 3). Cytochrome Oxidase I sequence data improved the resolution in identification of bacteria that use oxidative phosphorylation such as *Coxiella* and *Rickettsia*. These were mostly Rickettsia which could only be identified upto a genus level using 16S rRNA. 38 OTUs were classified into *Rickettsia* species, 16 OTUs were classified to *Rickettsia* subspecies and strains, and 7 OTUs were classified into *Coxiella burnetii* subspecies and strains.

**Table 3:**
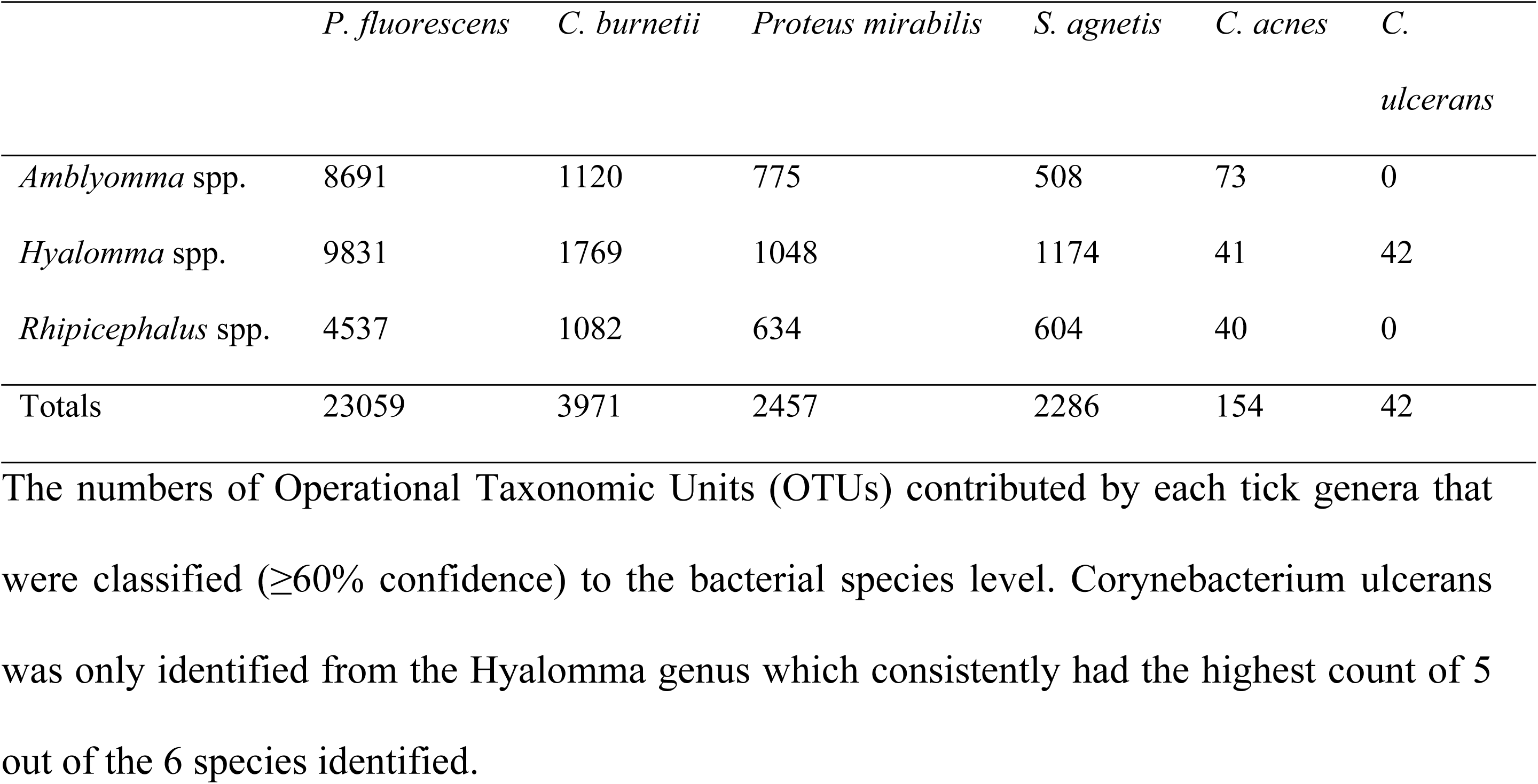
Distribution of OTUs that were classified to the bacterial species level per Tick Genera.

Consideration of bacterial sequences amplified using both 16S rRNA and COI approaches allowed for the identification of several pathogenic and potentially pathogenic bacterial species. These were *Coxiella burnetti*, *Proteus mirabilis*, *Cutibacterium acnes* and *Corynebaterium ulcerans*. Isiolo had a higher prevalence of pathogenic bacteria, particularly *Rickettsia conorii*, while Kwale’s leading pathogen was *Coxiella burnetii*. *Hyalomma truncatum* and *Rhipicephalus boophilus microplus* carried the most unique pathogens in Isiolo and Kwale, respectively.

COI sequence analysis on the BugSeq allowed for the speciation of the Rickettsia genus down to the species level. The platform identified 25 *Rickettsia* species of which 11 are known human pathogens: *Rickettsia parkeri, Rickettsia rickettsii, Rickettsia conorii, Rickettsia honei, Rickettsia slovaca Rickettsia japonica, Rickettsia sp Chad, Rickettsia aeschlimannii, Rickettsia helvetica, Rickettsia prowazekii, Rickettsia africae, Rickettsia vini,* and *Rickettsia massiliae. Rickettsia prowazekii* was the only Typhus Group (TG) *Rickettsia* that was identified. The pathogen was identified from *Amblyomma variegatum* ticks collected from Kwale county.

### Confirmation of Vector Identity using Cytochrome Oxidase Subunit I Sequences

COI sequence analysis confirmed the identities of all tick vectors reliably to the Genus level, however there were some discrepancies in 4.5% of the species between COI and morphological identification. Whereas these had been morphologically identified as *Rhipicephalus boophilus decoloratus* COI results identified them as *Rhipicephalus boophilus microplus.* The latter was used as the species identifier in this study.

## Discussion

Ticks are critical in both the biological and mechanical transmission of many pathogens especially when coupled with their ability to inhabit a wide variety of climates. Since ticks feed on multiple hosts throughout their lifecycles, it has become important under the One Health concept (3,30) to do constant surveillance of pathogens they carry to foster public health preparedness. This study sought to investigate the microbiomes of ticks and by extension the types of human pathogenic bacteria that are present in ticks collected from Kenya.

Ticks sampled in the present study were very skewed in favour of Isiolo (97.94%) against Kwale county (2.06%) (S1 Fig). The movement of livestock in Isiolo is higher at both inter and intra county levels (31) as compared to Kwale county which may also explain the higher number of tick infestation in Isiolo. Additionally, this disparity in collection may be because two of the sites from Isiolo were a livestock market and a slaughterhouse respectively, which would receive a disproportionate number of livestock and by extension ticks.

Previous entomological studies show that *Amblyomma variegatum* ticks prefer to feed on horses, cattle, sheep, dogs, and humans as hosts (32). *Hyalomma* species are known to preferentially feed on rabbits, hares, rodents, and birds while they are nymphs but switch to mostly ruminants once they become adults (33) whereas *Rhipicephalus boophilus microplus* has been shown to preferentially feed on cattle(34). The presence of *Cutibacterium acnes,* a bacterium that is commonly associated with human skin (35), in the ticks sampled in this study shows potential feeding on human hosts. *Amblyomma variegatum* ticks had the highest prevalence of *Cutibacterium acnes* which can be explained by their higher preference to feed on humans compared to the other species.

The alpha diversity test showed that richness and evenness varied at different taxonomic levels. However, the variability was not statistically significant (Table 1). This would suggest similar distribution of the bacterial communities among the different tick genera. Although the bacteriome in ticks is influenced by host, environmental and trans-ovarian factors (28), commensal or symbiotic relationships also drive the presence of some of these bacteria since they are useful or important for the survival of the ticks (36,37). These relationships between the ticks and endosymbiotic bacteria might explain why the overall bacteriome of different tick species collected from the two different regions is similar to a greater extent as indicated by the alpha diversity indices. This is probably because the presence of some of these endosymbionts increases the general survivability and fitness of the tick (36). The beta diversity test also indicated that the difference in bacterial abundances between the different genera, was not significant. In addition to this, the bacterial groups especially at the order level were somewhat similar to each other. Thus, bacterial infestation may be influenced by external factors other than the type of tick. These factors may include type of host, local environmental conditions such as temperature (38), or transmission between consecutive generations (39).

The present study identified Pseudomonadales, Rickettsiales and Coxiellales as the most abundant bacterial orders sampled in Isiolo and Kwale. The Pseudomonadales identified in this study include *Pseudomonas fluorescens* (Table 3), which is a ubiquitous bacterium in the environment that has recently been implicated as a potential pathogen (40). The majority of Rickettsiales identified in this study belonged to the genus Rickettsia (Fig 4). Prior studies screening ticks have demonstrated the abundant presence of Rickettsiae spp. in Kenya (40–42). Similarly, Rickettsial diseases such as tick typhus and African tick bite fever are well established in Kenya. This includes a recent fatal case of *Rickettsia conorii* infection in an American traveller in Kenya (41), and a *Rickettsia felis i*nfection that afflicted six people in the Northeastern region of Kenya (42). Rickettsial diseases have persisted in the region for over fifty years; however, there are still knowledge gaps in surveillance and reporting that need to be filled (43).

The most notable member of the Coxiellales identified in this study was *Coxiella burnetii* (Fig 4). These pathogenic intracellular bacteria are responsible for Q Fever, a disease commonly identified in people working close to farm animals or in pastoral environments (44). The high prevalence of this pathogen amongst ticks sampled from animals in the present study portends a risk to public health (Table 4). A previous study in Kenya has linked *C. burnetii* transmission in humans to ticks (45). Camels, in particular, have been identified as playing a bigger role in transmission of Q-fever compared to other ruminants (35). In the present study camels contributed 147 (8.9%) of the 954 ticks that had *C. burnetii* (Table 4). Whereas a recent study did not find a significant association between high prevalence of *Coxiella burnetii* in livestock and their owners (46), children have been shown to have a higher predisposition to infection as this pathogen can be transmitted through breast milk (47).

**Table 4:**
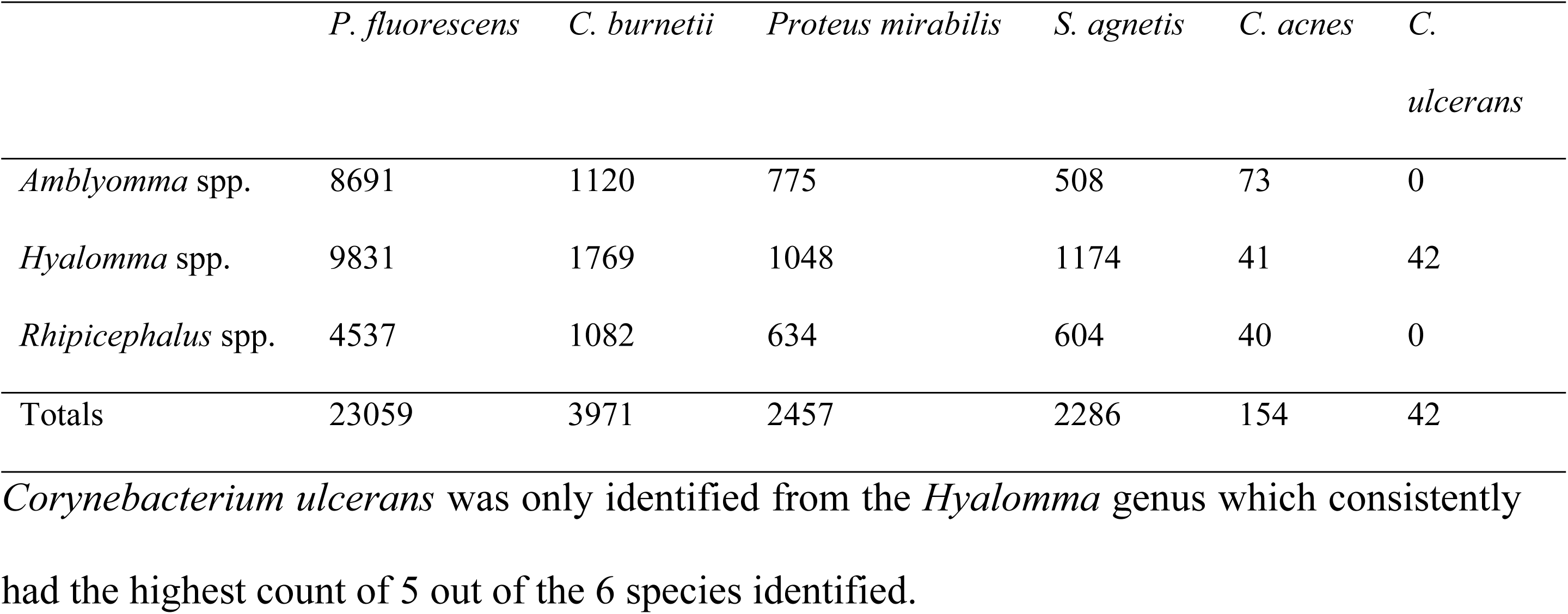
The numbers of Operational Taxonomic Units (OTUs) contributed by each tick genera that were classified (≥60% confidence) to the bacterial species level.

Comparative analysis of result from this and previous studies has highlighted that there are fluctuations in the detected prevalences of *C. burnetii*. On the lower end, one failed to detect it (48) and another reported a prevalence of 5.5% (49). On the higher end, a study reported a prevalence of 45.7% (50) which is comparable to the current detected level of 37.88%. This shows that more research is required to ascertain what other factors may be affecting the prevalence of *C. burnetii* in ticks. This could be, seasonal variations, county, hosts, sampling strategy, methods, and unknown outbreaks. This study included marketplaces and slaughterhouses as part of the sample sites as compared to the others that focused solely on farms (49,51). This study was unable to detect the presence of any *Anaplasma, Theileria* and *Borrelia* species detected in one of the previous studies(48), conducted on ticks collected from wildlife. This may be due to the ticks in the wildlife study having a wider range of vertebrate hosts each thus increasing the pool of potential reservoirs for the pathogens.

Analysis of tick species and the bacterial pathogens (Fig 5) indicates that all *Rhipicephalus pulchellus* pools had *Coxiella burnetii* and four Rickettsia species (*Rickettsia honei*, *Rickettsia japonica*, *Rickettsia sibrica* and *Rickettsia aeshlimannii*). This contrasts with an earlier study that reported a 25% prevalence of *Coxiella burnetii* within *Rhipicephalus* tick species (51), this may however be explained by the super-pools of ticks tested which had seventy-nine ticks. This means that testing of more ticks collected at different seasons (longitudinal study) and not oversampling individual animals may be required to ascertain other extraneous variables that may affect the prevalence of *Coxiella burnetii. Rickettsia prowazekii* was the only Typhus Group (TG) rickettsia detected. It was localized to *Amblyomma variegatum* species of ticks collected from Kwale county (Fig 5). The Typhus Group rickettsiae are commonly associated with fleas and body lice (52). *Rickettsia prowazekii* infection (epidemic typhus) has a reported fatality rate of 15% that goes even higher in regions of high poverty and low medical access and support (53) such as those in Low- and Middle-Income countries, including Kenya.

**Fig 5:**
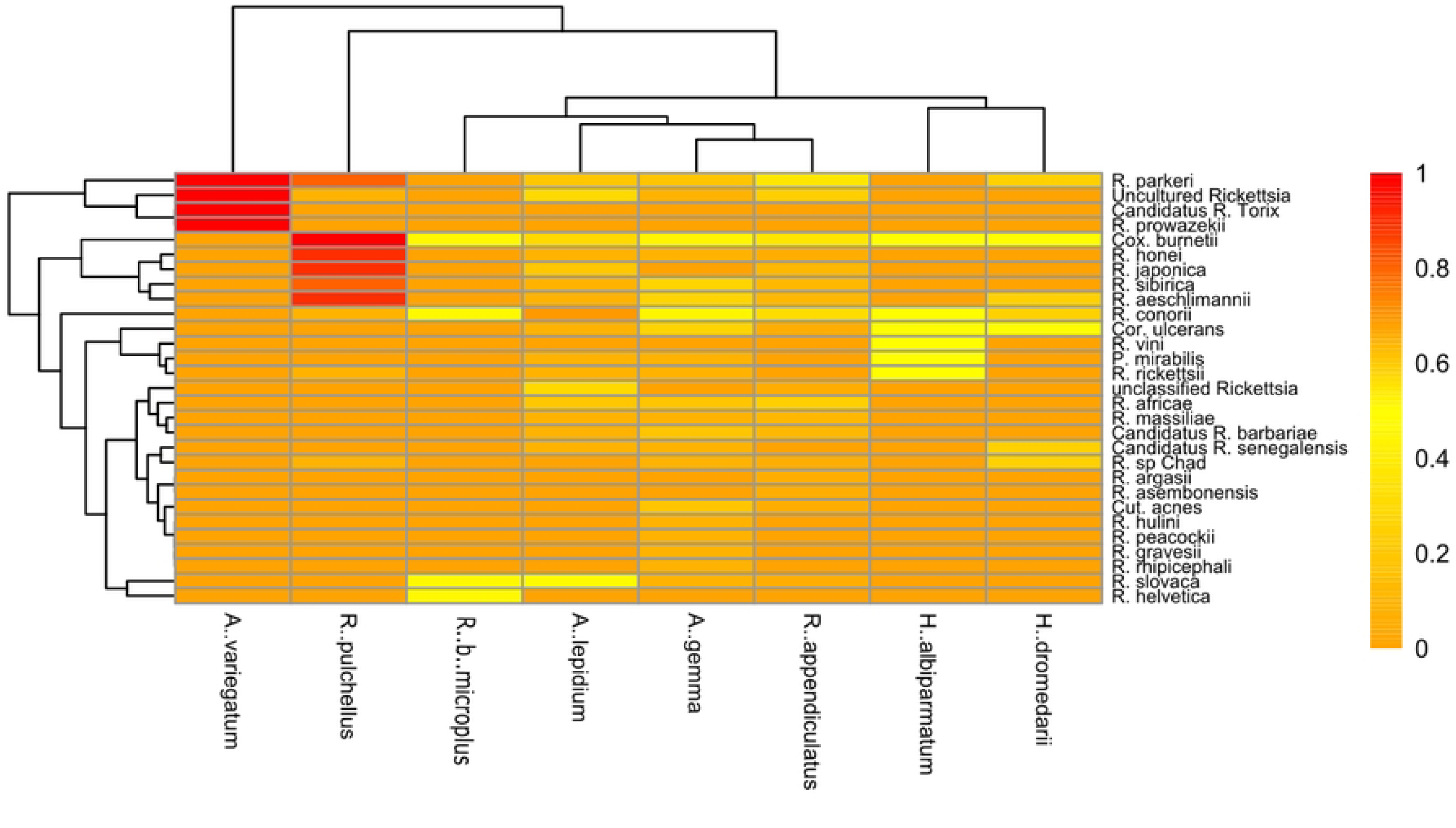
Heatmap showing the relative abundance of different bacterial pathogens identified. These include *Rickettsia* species identified using Cytochrome C Oxidase subunit 1 in tick species sampled.

Another contrast was seen with *Proteus mirabilis*, which was detected in lower levels than previous study (50). This difference may be due to the tick surface sterilization step in the current study, which removed external bacterial contaminants as *Proteus mirabilis* is widely found in soil and water.

This study has advanced the understanding of tick-associated bacteria especially with the combined use of 16S rRNA and COI sequence data to enhance the identification resolution for pathogenic bacteria to the species and strain levels for *Rickettsia* and *Coxiella* (54).

Our study, while illuminating, is not without limitations. Firstly, our sampling method, though rigorous, may not fully capture the entire tick population due to the potential localization of ticks in specific microhabitats. Moreover, our reliance on molecular techniques like 16S rRNA sequencing, while highly sensitive, carries inherent biases and risks of contamination. The lack of longitudinal data restricts our understanding of the temporal dynamics of tick-borne pathogens, that may be in flux depending on season. Additionally, our study’s focus on Isiolo and Kwale counties, while providing unique insights into these regions, limits the direct extrapolation of findings to areas with distinct environmental and socioeconomic contexts. Future research should address these limitations, potentially incorporating longitudinal studies across broader geographical regions. The impact of climate and seasonal variations during sampling should also be taken into consideration for a comprehensive understanding of tick-borne bacterial diseases in Kenya.

## Conclusion

In the current study, both intracellular and extracellular pathogenic bacteria known to cause human diseases were identified in ticks. Of interest is the high prevalence of *Coxiella burnetii* the causative agent of Q-fever and various pathogenic Rickettsia species identified in both Isiolo and Kwale, including *R. prowazekii* identified only in Kwale County. Lack of significant variation in microbiomes among the different tick species infers that ticks in the sampled areas share a core microbiome. This study has shown that analysis of full-length COI has the potential to be used to simultaneously identify bacteria that use oxidative phosphorylation for energy needs and their tick vectors to species level.

This study has highlighted the high prevalence of various pathogenic *Rickettsia* species and *Coxiella burnetti* in these regions in an effort to increase public health preparedness. The presence of *C. acnes* a bacterium commonly associated with human skin may be indicative of potential human feeding and exposure to the other pathogens. In light of this, there is a need to advise clinicians in these “at-risk” areas to include rickettsial diseases and query fever as part of their differential diagnosis. The viability of these bacteria and the competency of ticks to transmit them should be assessed in future studies.

## Acknowledgements

We thank Francis Ngere, Nicholas Odemba, David Oullo, Charles Waga, Richard Ochieng, Daniel Ngonga and Vitalice Opondo for their expert contribution in ticks sampling and identification.

This work was funded by the Armed Forces Health Surveillance Branch (AFHSB) and its Global Emerging Infections Surveillance (GEIS) Section, FY2022 ProMIS ID: P0116_22_KY and FY2023 ProMIS ID P0094_23_KY. The material has been reviewed by the Walter Reed Army Institute of Research. There is no objection to its presentation and/or publication. The opinions or assertions contained herein are the private views of the authors and are not to be construed as official or as reflecting the true views of the Department of the Army or the Department of Defence.

## Supporting Information

**S1 Fig: Percentage Stacked Bar Charts showing the abundances at the bacterial order level (a), genus level (b), and species level (c) per sample.** Samples with the’ ISL’ prefix are from Isiolo while those with the ’KWL’ prefix are from Kwale

**S1 Table: Distribution of ticks according to species and site.**

## References

1. Avise JC, Ayala FJ. From Wild Animals to Domestic Pets, an Evolutionary View of Domestication. National Academy of Sciences (US) [Internet]. 2009 [cited 2023 May 24]; Available from: https://www.ncbi.nlm.nih.gov/books/NBK219727/

2. Kaiser S, Hennessy MB, Sachser N. Domestication affects the structure, development and stability of biobehavioural profiles. Front Zool [Internet]. 2015 Aug 24 [cited 2023 May 24];12(Suppl 1). Available from: /pmc/articles/PMC5385816/

3. Aggarwal D, Ramachandran A. One Health Approach to Address Zoonotic Diseases. Indian J Community Med [Internet]. 2020 Mar 1 [cited 2023 Jun 29];45(Suppl 1):S6. Available from: /pmc/articles/PMC7232973/

4. Diaz JH. Introduction to Ectoparasitic Diseases. Mandell, Douglas, and Bennett’s Principles and Practice of Infectious Diseases. 2015 Jan 1;2:3243–3245.e1.

5. Rochlin I, Toledo A. Emerging tick-borne pathogens of public health importance: a mini-review. J Med Microbiol [Internet]. 2020 [cited 2023 May 29];69(6):781. Available from: /pmc/articles/PMC7451033/

6. Maxwell SP, Brooks C, McNeely CL, Thomas KC. Neurological Pain, Psychological Symptoms, and Diagnostic Struggles among Patients with Tick-Borne Diseases. Healthcare [Internet]. 2022 Jul 1 [cited 2023 May 29];10(7). Available from: /pmc/articles/PMC9323096/

7. Mac S, da Silva SR, Sander B. The economic burden of Lyme disease and the cost-effectiveness of Lyme disease interventions: A scoping review. PLoS One [Internet]. 2019 Jan 1 [cited 2023 May 29];14(1). Available from: /pmc/articles/PMC6319811/

8. Dong Y, Zhou G, Cao W, Xu X, Zhang Y, Ji Z, et al. Global seroprevalence and sociodemographic characteristics of Borrelia burgdorferi sensu lato in human populations: a systematic review and meta-analysis. BMJ Glob Health [Internet]. 2022 Jun 1 [cited 2023 May 29];7(6):e007744. Available from: https://gh.bmj.com/content/7/6/e007744

9. Ismail N, Bloch KC, McBride JW. Human Ehrlichiosis and Anaplasmosis. Clin Lab Med [Internet]. 2010 Mar [cited 2023 May 29];30(1):261. Available from: /pmc/articles/PMC2882064/

10. Chiuya T, Villinger J, Masiga DK, Ondifu DO, Murungi MK, Wambua L, et al. Molecular prevalence and risk factors associated with tick-borne pathogens in cattle in western Kenya. BMC Vet Res [Internet]. 2021 Dec 1 [cited 2023 May 29];17(1). Available from: /pmc/articles/PMC8627057/

11. Centers for Disease Control and Prevention. Imported Spotted Fevers [Internet]. 2019 [cited 2023 Oct 28]. Available from: https://www.cdc.gov/otherspottedfever/imported/index.html

12. Iruata MN, Wasonga OV, Ngugi RK, Kinuthia Ngugi R. Country Report Drylands and pastoralism Economic contribution of the pastoral meat trade in Isiolo County, Kenya Findings from Oldonyiro and Garbatulla Towns [Internet]. 2015. Available from: /theIIEDhttp://pubs.iied.org/10126IIED

13. Mohamed Sala S, Otieno DJ, Nzuma J, Mureithi SM. Determinants of pastoralists’ participation in commercial fodder markets for livelihood resilience in drylands of northern Kenya: Case of Isiolo. Pastoralism. 2020 Dec 20;10(1):18.

14. Kwale - Profile data - HURUmap Kenya: Making Census Data Easy to Use [Internet]. 2021 [cited 2021 Oct 10]. Available from: https://kenya.hurumap.org/profiles/county-2-kwale/

15. Okello-Onen J, Hassan SM, Suliman E. Taxonomy of African ticks: an identification manual. International Centre of Insect Physiology and Ecology (ICIPE); 1999.

16. Madder M, Horak I, Stoltsz H. Ticks: Tick identification. Pretoria; 2009.

17. Walker A, Bouattour A, Camicas JL, Estrada-Peña A, Horak I, Latif A, et al. Ticks of Domestic Animals in Africa: a guide to identification of species. 2003 Sep;

18. Paine GD. Ticks (Acari: Ixodoidea) in Botswana. Bull Entomol Res. 1982 Mar 10;72(1):1–16.

19. Matthysse JG, Colbo MH. The ixodid ticks of Uganda together with species pertinent to Uganda because of their present known distribution. College Park, MD: Entomological Society of America; 1987.

20. dos Santos HRM, Argolo CS, Argôlo-Filho RC, Loguercio LL. A 16S rDNA PCR-based theoretical to actual delta approach on culturable mock communities revealed severe losses of diversity information. BMC Microbiol. 2019 Dec 8;19(1):74.

21. Folmer O, Black M, Hoeh W, Lutz R, Vrijenhoek R. DNA primers for amplification of mitochondrial cytochrome c oxidase subunit I from diverse metazoan invertebrates. Mol Mar Biol Biotechnol. 1994 Oct;3(5):294–9.

22. Leger A, Leonardi T. pycoQC, interactive quality control for Oxford Nanopore Sequencing. J Open Source Softw. 2019 Feb 28;4(34):1236.

23. Ryan Wick. GitHub. 2018. Porechop, adapter trimmer for Oxford Nanopore reads.

24. Bolger AM, Lohse M, Usadel B. Trimmomatic: a flexible trimmer for Illumina sequence data. Bioinformatics. 2014 Aug 1;30(15):2114–20.

25. Bolyen E, Rideout JR, Dillon MR, Bokulich NA, Abnet CC, Al-Ghalith GA, et al. Reproducible, interactive, scalable and extensible microbiome data science using QIIME 2. Nat Biotechnol. 2019 Aug 24;37(8):852–7.

26. Rognes T, Flouri T, Nichols B, Quince C, Mahé F. VSEARCH: a versatile open source tool for metagenomics. PeerJ. 2016 Oct 18;4:e2584.

27. Bokulich NA, Kaehler BD, Rideout JR, Dillon M, Bolyen E, Knight R, et al. Optimizing taxonomic classification of marker-gene amplicon sequences with QIIME 2’s q2-feature-classifier plugin. Microbiome. 2018 Dec 17;6(1):90.

28. Leray M, Knowlton N, Machida RJ. MIDORI2: A collection of quality controlled, preformatted, and regularly updated reference databases for taxonomic assignment of eukaryotic mitochondrial sequences. Environmental DNA. 2022 Jul 11;4(4):894–907.

29. Wang Q, Garrity GM, Tiedje JM, Cole JR. Naïve Bayesian Classifier for Rapid Assignment of rRNA Sequences into the New Bacterial Taxonomy. Appl Environ Microbiol. 2007 Aug 15;73(16):5261–7.

30. Mackenzie JS, Jeggo M. The One Health Approach—Why Is It So Important? Trop Med Infect Dis [Internet]. 2019 May 31 [cited 2023 May 24];4(2). Available from: /pmc/articles/PMC6630404/

31. Iruata M, Wasonga O, Ngug R. Economic contribution of the pastoral meat trade in Isiolo County, Kenya: Findings from Oldonyiro and Garbatulla Towns. 2015.

32. Center for Food Security and Public Health. Amblyomma-variegatum. [cited 2023 Oct 29]; Available from: https://www.cfsph.iastate.edu/DiseaseInfo/notes/Amblyomma-variegatum.pdf

33. Spengler JR, Estrada-Peña A. Host preferences support the prominent role of Hyalomma ticks in the ecology of Crimean-Congo hemorrhagic fever. PLoS Negl Trop Dis. 2018 Feb 8;12(2):e0006248.

34. Tabor AE, Ali A, Rehman G, Rocha Garcia G, Zangirolamo AF, Malardo T, et al. Cattle Tick Rhipicephalus microplus-Host Interface: A Review of Resistant and Susceptible Host Responses. Front Cell Infect Microbiol. 2017 Dec 11;7.

35. Elston MJ, Dupaix JP, Opanova MI, Atkinson RE. Cutibacterium acnes (formerly Proprionibacterium acnes) and Shoulder Surgery. Hawaii J Health Soc Welf. 2019 Nov;78(11 Suppl 2):3–5.

36. Pollet T, Sprong H, Lejal E, Krawczyk AI, Moutailler S, Cosson JF, et al. The scale affects our view on the identification and distribution of microbial communities in ticks. Parasit Vectors. 2020 Dec 21;13(1):36.

37. Hussain S, Perveen N, Hussain A, Song B, Aziz MU, Zeb J, et al. The Symbiotic Continuum Within Ticks: Opportunities for Disease Control. Front Microbiol. 2022 Mar 17;13.

38. Couret J, Schofield S, Narasimhan S. The environment, the tick, and the pathogen – It is an ensemble. Front Cell Infect Microbiol. 2022 Nov 2;12.

39. Bonnet SI, Binetruy F, Hernández-Jarguín AM, Duron O. The Tick Microbiome: Why Non-pathogenic Microorganisms Matter in Tick Biology and Pathogen Transmission. Front Cell Infect Microbiol. 2017 Jun 8;7.

40. Liu X, Xiang L, Yin Y, Li H, Ma D, Qu Y. Pneumonia caused by Pseudomonas fluorescens: a case report. BMC Pulm Med. 2021 Dec 5;21(1):212.

41. Rutherford JS, Macaluso K, Smith N, Zaki SR, Paddock CD, Davis J, et al. Fatal Spotted Fever Rickettsiosis, Kenya. Emerg Infect Dis. 2004 May;10(5):910–3.

42. Richards AL, Jiang J, Omulo S, Dare R, Abdirahman K, Ali A, et al. Human Infection with *Rickettsia felis*, Kenya. Emerg Infect Dis. 2010 Jul;16(7):1081–6.

43. Richards A. Rickettsial diseases in Eastern Africa. International Journal of Infectious Diseases. 2014 Apr;21:66.

44. Dragan AL, Voth DE. Coxiella burnetii: international pathogen of mystery. Microbes Infect. 2020 Apr;22(3):100–10.

45. Knobel DL, Maina AN, Cutler SJ, Ogola E, Feikin DR, Junghae M, et al. Coxiella burnetii in Humans, Domestic Ruminants, and Ticks in Rural Western Kenya. Am J Trop Med Hyg. 2013 Mar 6;88(3):513–8.

46. Muema J, Nyamai M, Wheelhouse N, Njuguna J, Jost C, Oyugi J, et al. Endemicity of Coxiella burnetii infection among people and their livestock in pastoral communities in northern Kenya. Heliyon. 2022 Oct;8(10):e11133.

47. Kobbe R, Kramme S, Kreuels B, Adjei S, Kreuzberg C, Panning M, et al. Q Fever in Young Children, Ghana. Emerg Infect Dis. 2008 Feb;14(2):344–6.

48. Wang’ang’a Oundo J, Villinger J, Jeneby M, Ong’amo G, Otiende MY, Makhulu EE, et al. Pathogens, endosymbionts, and blood-meal sources of host-seeking ticks in the fast-changing Maasai Mara wildlife ecosystem. Telford III SR, editor. PLoS One [Internet]. 2020 Aug 1 [cited 2023 May 29];15(8):e0228366. Available from: /pmc/articles/PMC7458302/

49. Koka H, Sang R, Kutima HL, Musila L. Coxiella burnetii Detected in Tick Samples from Pastoral Communities in Kenya. Biomed Res Int. 2018;2018.

50. Ergunay K, Mutinda M, Bourke B, Justi SA, Caicedo-Quiroga L, Kamau J, et al. Metagenomic Investigation of Ticks From Kenyan Wildlife Reveals Diverse Microbial Pathogens and New Country Pathogen Records. Front Microbiol. 2022 Jul 1;13:2242.

51. Kumsa B, Socolovschi C, Almeras L, Raoult D, Parola P. Occurrence and Genotyping of Coxiella burnetii in Ixodid Ticks in Oromia, Ethiopia. Am J Trop Med Hyg [Internet]. 2015 Nov 4 [cited 2023 Mar 4];93(5):1074–81. Available from: https://www.ajtmh.org/view/journals/tpmd/93/5/article-p1074.xml

52. Onyiche TE, Labruna MB, Saito TB. Unraveling the epidemiological relationship between ticks and rickettsial infection in Africa. Frontiers in Tropical Diseases. 2022 Sep 6;3.

53. Sahni A, Fang R, Sahni SK, Walker DH. Pathogenesis of Rickettsial Diseases: Pathogenic and Immune Mechanisms of an Endotheliotropic Infection. Annual Review of Pathology: Mechanisms of Disease. 2019 Jan 24;14(1):127–52.

54. Clarridge JE. Impact of 16S rRNA gene sequence analysis for identification of bacteria on clinical microbiology and infectious diseases. Clin Microbiol Rev. 2004 Oct;17(4):840–62, table of contents.

